# Modeling the organizational heterogeneity of lipid-enriched microdomains in the neuronal membranes of gray and white matter of Alzheimer’s brain: A computational lipidomics study

**DOI:** 10.64898/2026.02.18.706545

**Authors:** Sruthi Peesapati, Sandipan Chakraborty

## Abstract

Alzheimer’s disease (AD) is a leading cause of death among the elderly, with no existing treatment. The development of therapy is further challenged by a limited understanding of molecular pathogenesis and the absence of reliable early-detection biomarkers. Neuroimaging and lipidomic studies reveal structural and biochemical alterations in both gray and white matter in AD patients, including disruptions in membrane organization and neuronal signaling pathways. In the present work, we employed lipidomics-guided modeling of membranes in gray and white matter regions in healthy and diseased (AD) conditions, and used all-atom molecular dynamics (MD) simulations to examine how AD-associated alterations in lipid composition influence the structure, spatial organization, and micro-heterogeneity of neuronal plasma membranes in different brain regions. Data suggest that Alzheimer’s disease–associated lipid alterations in gray matter (GM) and white matter (WM) impact membrane thickness and microdomain distribution, highlighting the critical role of lipid composition in maintaining neuronal membrane homeostasis and function. Higher-order cholesterol–ceramide– sphingomyelin–enriched domains are more abundant in the neuronal membranes of the GM region in diseased conditions. Under AD-mimicking conditions, lipidomic analyses demonstrate that neuronal membranes in GM experience more substantial compositional and structural remodeling than those in WM. Our results show significant changes in membrane microdomain distribution across the lipid bilayers, and, interestingly, these changes are more pronounced in the gray matter than in the white matter. This study establishes a framework for modeling the tissue-specific lipidomics data to understand how disease-driven compositional changes affect the structure, organization, and dynamics of biological membranes.

## Introduction

Alzheimer’s disease (AD) is a chronic and progressive neurodegenerative disorder and is a leading cause of dementia-related death worldwide. Clinically, it manifests with gradual memory impairment, decline in cognitive functions, and loss of functional independence^1,2^. Pathologically, AD is characterized by the accumulation of amyloid-β (Aβ) plaques, intracellular deposition of neurofibrillary tangles composed of hyperphosphorylated tau, and synaptic dysfunction^3–5^. These alterations are considered to be a critical factor contributing to the irreversible cognitive decline observed among AD patients, and are accompanied by progressive brain atrophy. Despite extensive efforts, the current medications for AD remain largely palliative, providing transient symptomatic relief without halting disease progression or restoring cognitive functions. Although active research is ongoing to explore different therapeutic modalities, including natural and synthetic compounds, peptide and peptidomimetic agents, and amyloid-targeting immunotherapies^6–16^, their success in clinical trials remains very limited. Moreover, these therapies are associated with side effects, underscoring the need for effective disease-management strategies. A major obstacle in the management of AD is delayed diagnosis, as detectable cognitive deficits appear only at advanced stages of neurodegeneration. Consequently, identifying reliable biomarkers for capturing pathological changes remains challenging.

Neuroimaging studies of patients with AD have shown structural and biochemical alterations in both gray matter (GM) and white matter (WM)^17,18^. Specifically, GM, which is enriched with neuronal cell bodies and synaptic membranes, undergoes severe atrophy due to neuronal loss^19^. Patients with AD reportedly exhibited reductions in inner gray matter volume and whole brain volume^20–23^. In contrast, WM, which is composed predominantly of myelinated axonal membranes, shows demyelination, axonal degeneration, and reduced cholesterol content in AD patients^24,25^. These changes indicate disrupted myelin integrity and altered membrane composition. Neuroimaging and morphometric analyses have revealed early GM loss confined to medial temporal regions that later extends to parietal and frontal lobes, along with concomitant WM abnormalities, indicating impaired neural connectivity^26^. Studies on human postmortem brain samples and appropriately developed mouse models have shown region-specific lipid dysregulation^27,28^.

Growing evidence indicates that dysregulation of lipid metabolism is a key feature of AD pathology^29–42^. Multiple studies have documented alterations in phospholipids (phosphatidylcholine, phosphatidylethanolamine, phosphatidylinositol, and phosphatidylserine), sphingolipids (sphingomyelin, cerebrosides, ceramides, sulfatides, and gangliosides), and cholesterol composition in AD brains. It is widely accepted that lipids play a central role in maintaining the structural and functional integrity of neuronal membrane architecture. In addition, lipids play a critical role in intracellular signaling in brain^43–45^. Any deviations in lipid composition would alter interfacial properties, such as membrane thickness, spatial organization, and curvature, and would further influence interactions with proteins and signaling pathways^29^. The complexity is further compounded by the asymmetric distribution of lipids across the bilayer leaflets, a feature whose functional advantages and biological significance require further investigation. Studies have also shown that more lipids are differentially expressed in GM than in WM^37^.

Despite the growing recognition of lipid dysregulation in AD, the molecular-level mechanisms by which lipid compositional perturbations influence the membrane structure and organization of different neuronal cells in GM and WM remain poorly understood. In this study, we employed atomistic MD simulations to investigate alterations in the biophysical properties of neuronal membranes in different brain regions (GM and WM) under AD conditions, modelled using computational lipidomics. By systematically comparing membrane structural properties, lipid organization, and dynamics, we aim to elucidate how disease-associated lipid perturbations modulate membrane architecture across distinct brain regions. These findings provide molecular-level insight into the lipid-driven mechanisms that may underlie the differential vulnerability of GM and WM to AD pathology.

## Methodology

All simulations were performed using the GROMACS 2020.6 package^46–48^. The all-atom CHARMM36 force field was used to parameterize the lipid systems^49^. Healthy and diseased lipid bilayer membranes were constructed using the membrane modeler module of CHARMM-GUI^50^. The lipid composition was modelled based on the experimental reports of Lam *et al.*^30^ Membrane asymmetry was incorporated based on the published literatures^51,52^. Lipid composition of each membrane in different conditions is summarized in Table S1. The initial box dimensions were 22 × 22 × 8.5 nm³. Each bilayer was placed in the center of the simulation box and immersed in TIP3P water. After complete solvation, the water molecules within the lipid bilayer’s hydrophobic core were removed using a Perl script. This was followed by the addition of Na^+^ and Cl^-^ ions to maintain a 150 mM salt concentration for all the systems. Subsequently, all systems were energy-minimized using the steepest descent algorithm. Then, each system was subjected to a series of equilibration simulations comprising a 5 ns simulation in the NPT ensemble (1 fs time step) and a 5 ns simulation in the NVT ensemble (2 fs time step), followed by 10 ns of equilibration in the NPT ensemble (2 fs time step). The production runs were performed in an NPT ensemble for 500 ns (2-fs time step) for each system. This procedure was repeated in triplicate for each system, with independent initial velocity distributions. Temperature and pressure were controlled using a Noose-Hoover thermostat and a semi-isotropic Parinello-Rahman barostat, with coupling constants of 0.5 ps and 5 ps, respectively. The particle mesh Ewald (PME) summation technique was used with a grid spacing of 0.12 nm.

### Analysis tools

The Visual Molecular Dynamics package (VMD) was used to visualize the simulation trajectories^53^. Built-in GROMACS tools have been used for trajectory post-processing and preliminary analysis. FATSLiM was used to determine the area per lipid and thickness of the membrane bilayers^54^. The 2D membrane thickness was calculated using MEMBPLUGIN in VMD^55^. Custom Python code was written and used to determine microdomain sizes, their corresponding counts, and residence times.

## Results and Discussion

Biological membranes are known for their intricate complexity in composition, asymmetry, and spatiotemporal organization. The brain is the second most lipid-rich organ after adipose tissue, with lipids constituting over 50 percent of its dry weight^56^. This highlights the importance of structural lipids in maintaining neuronal membrane integrity. Here, using multiple all-atom MD simulations, we have studied variations in the structural and spatial organization of lipids in the neuronal membranes in the GM and WM regions, appropriately modelled from lipidomics data from postmortem frozen brain tissues of subjects with AD pathology, namely Braak stage V-VI, and moderate-severe vascular pathology (referred as ADB system in the manuscript), and compared with age-matched controls (referred as HB system in the manuscript). A total of 9 lipids, including cholesterol (CHL), 1-palmitoyl-2-oleoyl-*sn*-glycero-3- phosphocholine (POPC), 1-palmitoyl-2-oleoyl-*sn*-glycero-3-phosphoethanolamine (POPE), 1- palmitoyl-2-oleoyl-*sn*-glycero-3-phosphoserine (POPS), 1-palmitoyl-2-oleoyl-*sn*-glycero-3- phosphoinositol (POPI), *N*-palmitoyl-d-erythro-sphingosinephosphorylcholine (PSM), *N*- palmitoyl-d-erythro-sphingosine (CER), 1-palmitoyl-2-oleoyl-sn-glycero-3-phosphatidic acid (POPA), 1-Palmitoyl-2-oleoylphosphatidylglycerol (POPG) are part of the membrane model, representing the major components of neuronal membranes. Figure 1A shows the compositional heterogeneity of the outer leaflet (OL) and inner leaflet (IL) of the four systems studied in this work, with each lipid colored uniquely.

**Figure 1:**
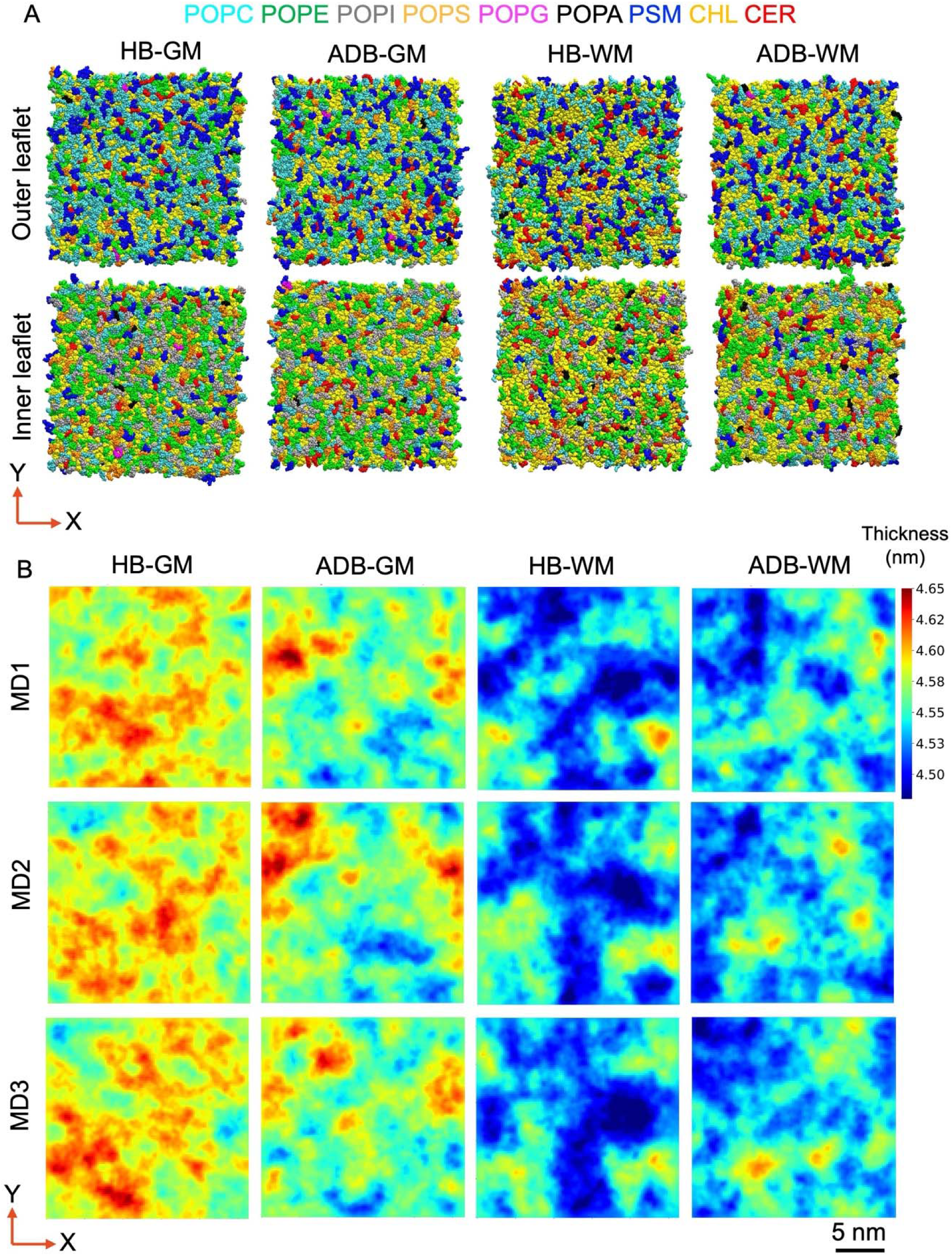
A) Schematic representation of outer and inner leaflets showing the spatial distribution of lipid components in the GM and WM regions modelled from the lipidomics data of postmortem frozen brain tissues of subjects with AD pathology with moderate-severe vascular pathology, and healthy age-matched control, with each lipid colored uniquely. B) 2D membrane thickness as determined using the Phosphate atoms of phospholipids (POPC, POPE, POPI, POPS, POPA, POPG) and sphingolipids (PSM) as the reference by averaging over 0.5 μs of the simulation trajectories. (HB: healthy membrane, ADB: AD-mimicking diseased model membrane, GM: gray matter, WM: white matter).

### Effect of altered lipid composition in diseased conditions on the structural properties of membranes

Figure S1 displays the number density profiles of all membrane lipid components for each system studied in this work. These profiles clearly reflect the compositional differences between the membranes, as the density distributions of each species directly correspond to their relative abundances in the bilayer. Across both neuronal membrane models of the GM and WM regions, the mean area per lipid (APL) did not differ significantly between healthy (HB) and diseased (ADB) membranes (Table S2). However, compared with the neuronal plasma membranes in the GM, the axonal membranes of the WM region consistently showed a slightly inner APL. Membrane thickness (Table S2) of the neuronal plasma membranes of the GM region exhibited a decrease of 0.02 nm in the diseased condition as compared to healthy membranes. However, changes in membrane thickness are insignificant in axonal neurons of the WM region in both healthy and diseased conditions. To further understand the spatial variation of bilayer thickness across the bilayer plane, we generated two-dimensional (2D) thickness maps. Figure 1B shows the representative 2D thickness profiles for all the systems. In GM, the healthy membrane exhibits homogeneous regions of higher thickness (> 4.60 nm), whereas in diseased conditions, the membrane displays a more heterogeneous pattern with localized regions of both higher and lower thickness (< 4.53 nm). In the WM region, healthy membranes exhibit localized regions of low overall thickness. In AD, there is an increase in the regions with high thickness. However, the changes are less pronounced than those observed in the membranes of the GM region, in both healthy and diseased conditions. With the change in membrane composition, the overall thickness profiles vary, with no major change in interdigitation across all systems. Furthermore, the mean cholesterol tilt angle decreases in the diseased membrane compared to the healthy membrane in the GM region, suggesting that a greater proportion of cholesterol molecules are strongly aligned with the bilayer normal, indicating preferential engagement in higher-order organization.

### Effect of altered lipid composition on the spatial distribution of the lipids: Enrichment analysis

The radial distribution function (RDF) profile of cholesterol ‘O3’ atoms around cholesterol ‘O3’ atoms showed two distinct peaks at 0.57 nm and 1.08 nm for all four systems studied (Figure S2). Similarly, the radial distribution of ceramide and sphingomyelin around cholesterol showed a minor peak at 0.28 nm and a higher-intensity peak at 0.5 nm. Furthermore, to investigate the spatial organization of membrane lipids, we have examined the enrichment profiles of each lipid relative to the others in both healthy and diseased membrane models of the GM and WM regions. We have calculated lipid enrichment profiles using a cutoff distance of 0.88 nm (the entire 1^st^ peak of the RDF profiles). Lipid enrichment values quantify the extent to which a particular lipid is preferentially organized around a reference lipid, with values above 1 indicating enrichment and those below 1 indicating depletion. Figure 2A & 2B show the average lipid enrichment heatmap of all lipids in the lipid bilayer, specifically leaflet-wise, for healthy (HB) and diseased (ADB) model membranes in the GM and WM regions. Figures S3 and S4 show the distribution profile of specific lipids relative to cholesterol and ceramide, respectively.

**Figure 2:**
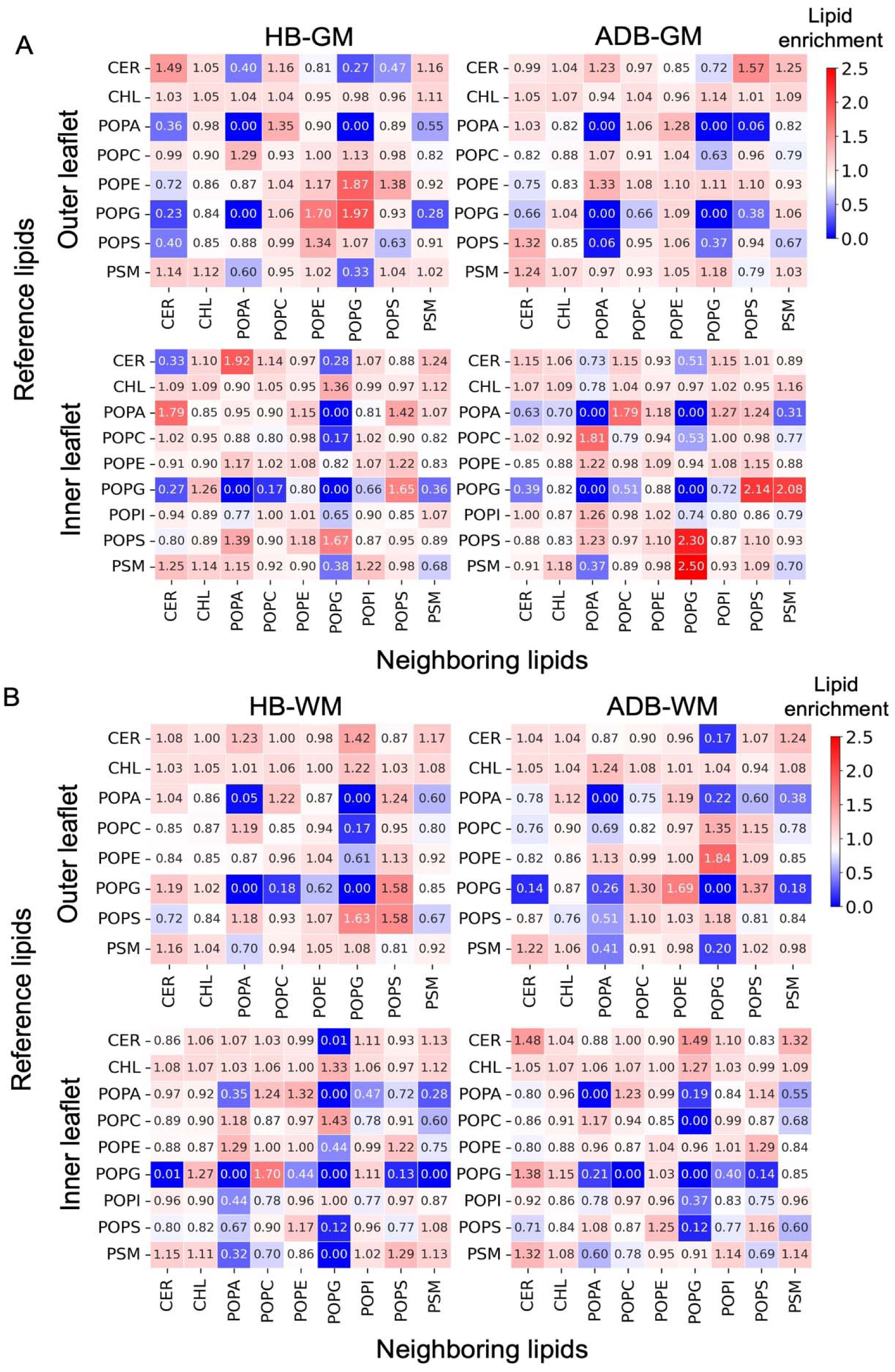
A) Lipid enrichment index heat map averaged over the simulation timescale for the outer leaflet and inner leaflet of healthy and diseased membranes in the GM region. B) Lipid enrichment heat map averaged over the simulation timescale for the outer leaflet and inner leaflet of healthy and diseased membranes in the WM region. The data are averages over the concatenated 1.5 μs trajectory for each system. Hydroxyl oxygen atom (‘O3’) is considered a reference for ceramide and cholesterol, while Phosphate atoms are considered a reference for the rest of the lipids. (HB: healthy membrane, ADB: AD-mimicking diseased model membrane, GM: gray matter, WM: white matter).

Tables S3 and S4 show the corresponding “area under the curve” of enriched regions (the enrichment index ≥ 1). A minor increase in cholesterol self-enrichment is observed only in the outer leaflet of the membranes that belong to the GM region in the AD condition. Ceramide enrichment around cholesterol increases in the outer leaflet of the membranes in the GM as well as WM region in the disease-mimicking membrane composition. Notably, both cholesterol and ceramide concentrations increase in the GM region in diseased conditions. Although the ceramide and cholesterol concentrations are uniform across the leaflets and vary only among systems, the enrichment profiles of ceramide and cholesterol are markedly different between the leaflets in the GM region in healthy as well as diseased conditions. Sphingomyelin enrichment around cholesterol remains almost similar in the outer leaflet of healthy and diseased membrane models of GM and WM regions.

### Characterization of Cholesterol-Ceramide-Sphingomyelin-enriched functional microdomains in membranes of different brain regions and their changes in diseased conditions

The lateral distribution of lipids within biological membranes is heterogeneous, with specific lipids self-organizing to form distinct microdomains. These functional microdomains are believed to be largely organized by the association of cholesterol, sphingomyelin, and ceramide, which form the platform for cellular signaling. We have evaluated the spatial organization of these microdomains by calculating their 2D number density distributions across all replicas of each system. Figure 3A illustrates the schematic representation of the microdomain number density with the density gradient ranging from blue (low) to red (high). In the GM region, the contrast in density is visually evident for both the outer and inner leaflets of the diseased membranes. On the other hand, for the WM region, such differences are less apparent. We have further classified and quantified the number density (coded into color density) into five density regions (Figure 3B): low (L: 0-0.1), intermediate (I: 0.1-0.2), medium1 (M1: 0.2-0.3), medium2 (M2: 0.3-0.4), and high (H: 0.4 and above). In the GM region, microdomains with a low number density (L) are abundant in the inner leaflet, particularly in healthy membranes. In the WM region, microdomains of low number density are abundant in the inner leaflet, but no significant differences are observed between healthy (HB) and diseased (ADB) conditions in both leaflets. The microdomains with intermediate density (I) remain similar for the outer and inner leaflets of the membranes in the GM region, but they are higher in the inner leaflet of the WM region (Figure 3B). The microdomains corresponding to the medium-density (M1 and M2) regions show increased abundance in diseased membranes, both in the outer and inner leaflets in the GM region. Notably, for GM membranes, large, high-density microdomains are relatively rare compared to low-density microdomains. regions. In the case of WM, the most abundant microdomains are primarily low-, intermediate-, and medium-density, and there is almost no difference in their numbers between healthy and diseased conditions.

**Figure 3:**
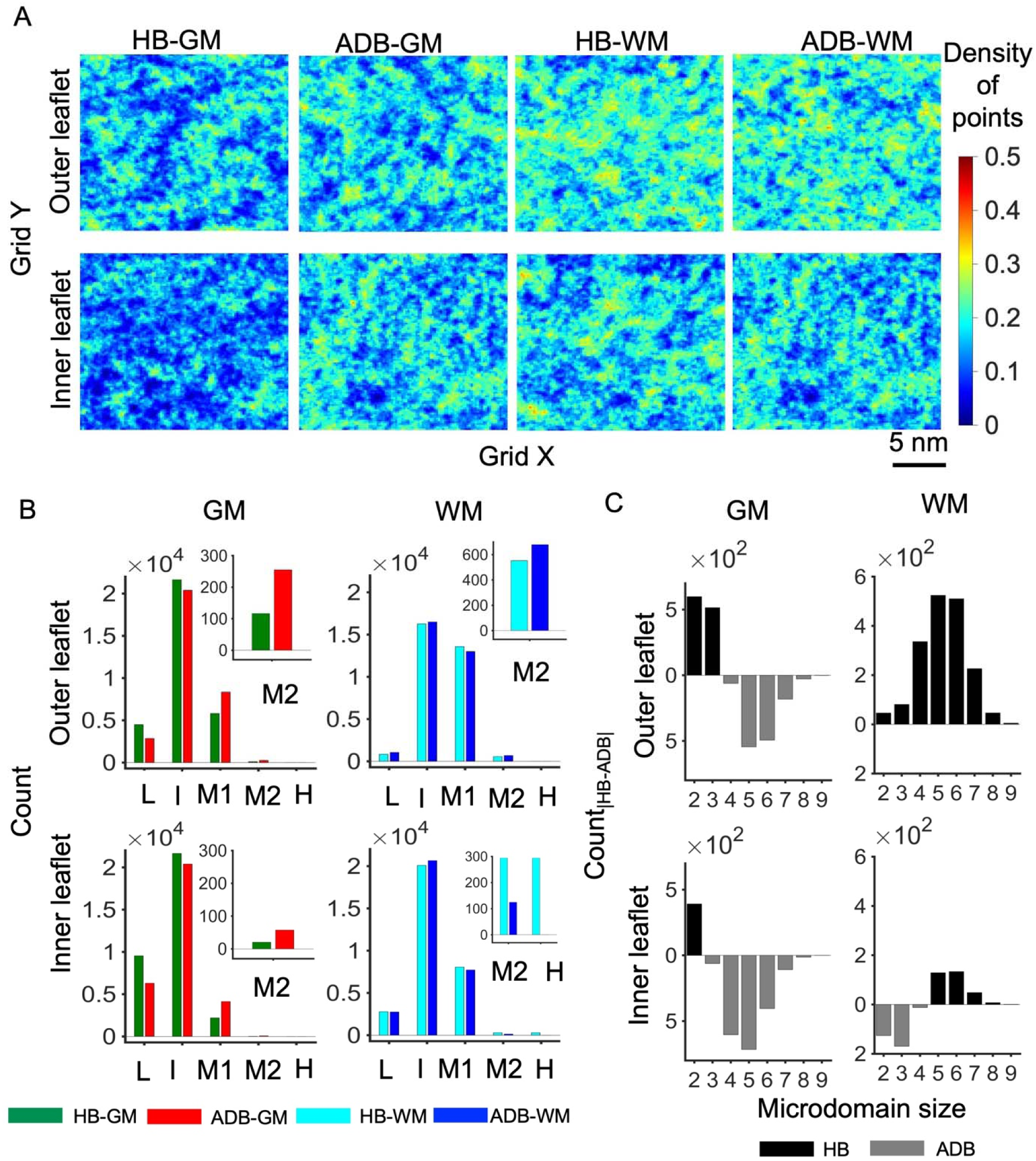
A) 2D number density distribution of cholesterol, ceramide, and sphingomyelin using the hydroxyl oxygen atoms (‘O3’) averaged over the entire simulation trajectories for all the systems. B) Qualitative analysis of 2D number density profiles, categorized into five distinct regions, based on the intensity profiles of the number density: low (L), intermediate (I), medium1 (M1), medium2 (M2), and high (H) corresponding to 0-0.1, 0.1-0.2, 0.2-0.3, 0.3-0.4, and 0.4-0.5 number density values, respectively. C) Size distributions of microdomains in each leaflet have been obtained by calculating the near-neighbor aggregates (using a cut-off of 8.8Å) from simulation trajectories. The difference in the number of aggregates formed by cholesterol, ceramide, and sphingomyelin in the outer and inner membrane leaflets in the GM and WM regions, under healthy and AD conditions, is shown. The data are normalized by the number of cholesterol molecules. (HB: healthy membrane, ADB: AD-mimicking diseased model membrane, GM: gray matter, WM: white matter).

We have further characterized the size of microdomains in each leaflet of all four systems by calculating the near-neighbor aggregates from simulation trajectories. Figure 3C shows the difference in the count of microdomains of varying sizes between healthy and diseased conditions. Clearly, the microdomains of higher order (size 4 and above) are abundant in diseased conditions (ADB) for both the membrane leaflets in the GM region, while changes are less significant for membranes in the WM region. Thus, in AD conditions, there is increased cholesterol-mediated aggregation in the neuronal plasma membranes of the gray matter region.

### Comparative changes in microdomain compositions and their stability in membranes of different brain regions in healthy and ADB conditions

We have further elucidated the compositional variations of microdomains with aggregation numbers ranging from 3 to 8 in healthy and diseased membrane models of GM and WM regions. Furthermore, we have also determined the residence time (the duration during which the microdomain components remain together within the cutoff distance before diffusing away) of each composition of a particular microdomain size (Figure S5-S10). The counts for each possible combination are compared across leaflets for healthy and diseased conditions. For microdomains of size 3 (Figure 4), the cholesterol-only domains are abundant in the inner leaflet than in the outer leaflet in all the studied conditions. In particular, the number of cholesterol-only microdomains in membranes is higher in the GM than in the WM regions in both healthy and disease-mimicking conditions. The number of domains with 2 ceramides and 1 cholesterol is abundant and stable for the inner leaflet in the diseased membrane of the GM regions (Figure S5).

**Figure 4:**
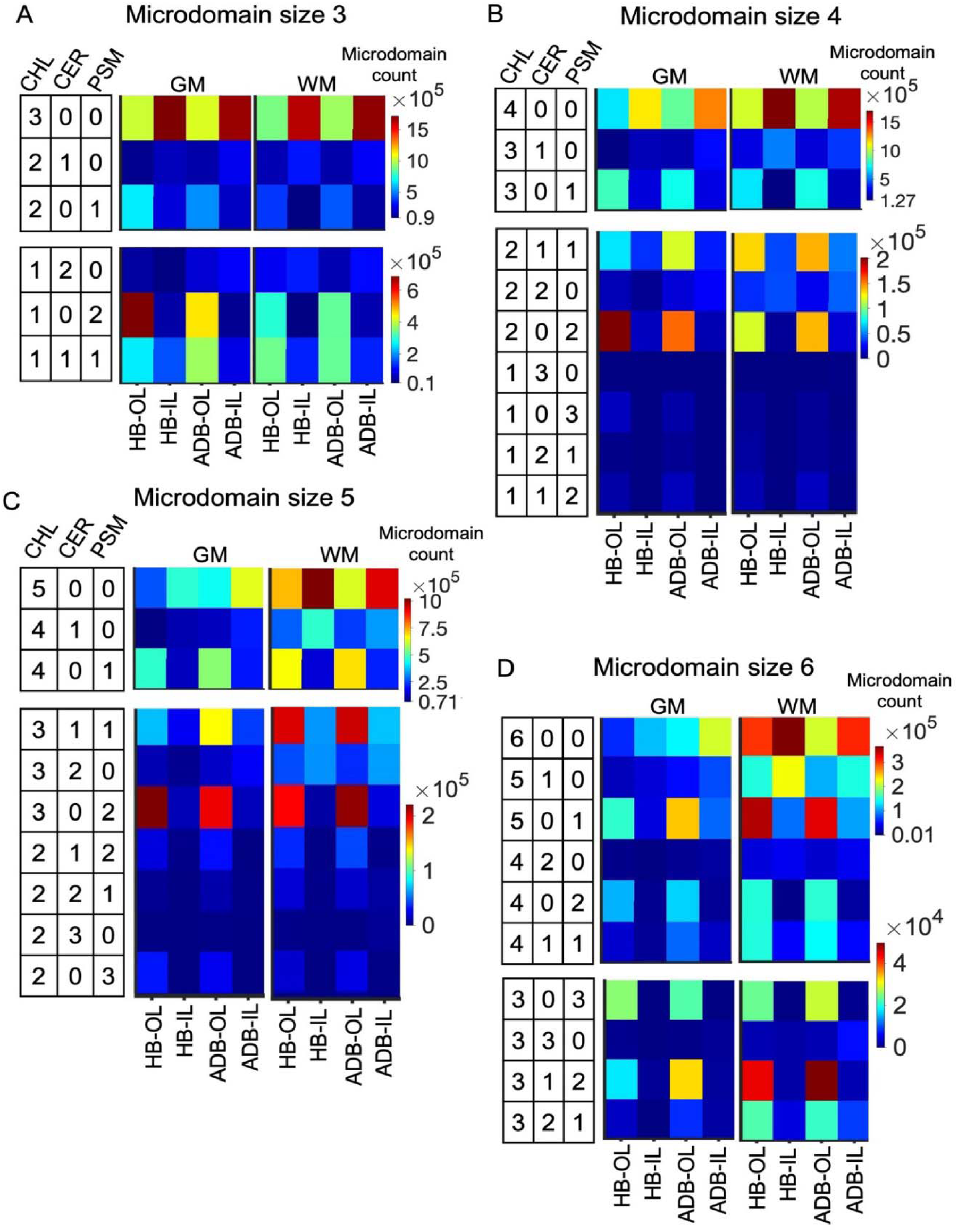
A) Compositions of microdomains of size 3 and their corresponding counts as obtained from the 1.5 μs simulation trajectory of healthy and diseased membranes, including both leaflets, of GM and WM regions. B) Compositions of microdomains of size 4 and their corresponding counts as obtained from the 1.5 μs simulation trajectory of healthy and diseased membranes, including both leaflets, of GM and WM regions. C) Compositions of microdomains of size 5 and their corresponding counts as obtained from the 1.5 μs simulation trajectory of healthy and diseased membranes, including both leaflets, of GM and WM regions. D) Compositions of microdomains of size 6 and their corresponding counts as obtained from the 1.5 μs simulation trajectory of healthy and diseased membranes, including both leaflets, of GM and WM regions. (HB: healthy membrane, ADB: AD-mimicking diseased model membrane, OL: outer leaflet, IL: inner leaflet, GM: gray matter, WM: white matter).

For microdomains of size 4 (Figure 4), inner leaflets are more enriched with cholesterol-only microdomains in the membranes of the GM and WM regions. Interestingly, in the GM region, cholesterol-only microdomains are more abundant in diseased conditions. Although their stability remains similar in the healthy and diseased conditions (Figure S6, Supporting Information). Interestingly, microdomains containing one or more ceramides and cholesterol are abundant and stable in diseased membranes. Similarly, sphingomyelin-enriched microdomains (without ceramides) are abundant in the healthy brain membrane, particularly in the outer leaflets (Figure S6, Supporting Information). For microdomains of size 5 & 6 (Figure 4), Cholesterol-only and cholesterol-enriched microdomains are more abundant in the diseased membrane in comparison to the healthy membrane models of gray matter. However, this is not the case with white matter, where there is a decrease of highly cholesterol-enriched microdomains in the diseased condition. Ceramide-enriched and sphingomyelin-depleted microdomains are more evident, and they are stable in the diseased model membrane of the GM region (Figure S7 & S8, Supporting Information).

For microdomains of size 7 and 8 (Figure 5), the cholesterol-only and primarily cholesterol-enriched microdomains are more prevalent in the membranes of gray matter in diseased conditions. The observation is more pronounced when the microdomains are composed of higher cholesterol content, with 1 or 2 sphingomyelins, but are devoid of ceramides. However, in white matter, cholesterol-only and primarily cholesterol-enriched microdomains are either prevalent in healthy membranes or are not significantly different between healthy and diseased membranes. Interestingly, cholesterol-only and primarily cholesterol-enriched microdomains are more stable in the gray matter of the diseased membrane model (Figure S9 & S10, Supporting Information).

**Figure 5:**
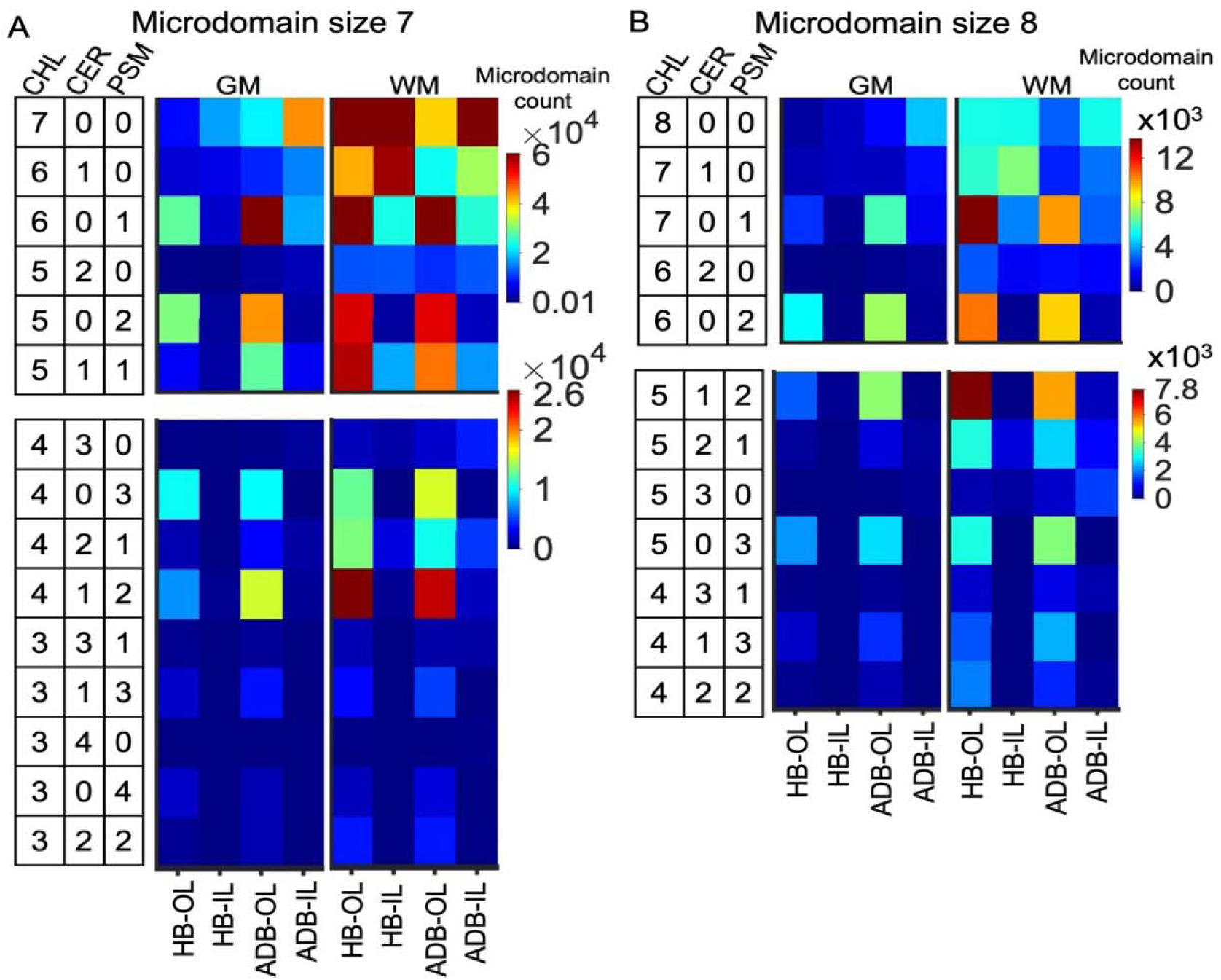
A) Composition of microdomains of size 7 and their corresponding counts as obtained from the 1.5 μs simulation trajectory of healthy and diseased membranes, including both leaflets, of GM and WM regions. B) Composition of microdomains of size 8 and their corresponding counts as obtained from the 1.5 μs simulation trajectory of healthy and diseased membranes, including both leaflets, of GM and WM regions. (HB: healthy membrane, ADB: AD-mimicking diseased model membrane, OL: outer leaflet, IL: inner leaflet, GM: gray matter, WM: white matter).

### Influence of POPI in regulating the microdomain distributions in diseased conditions

Phosphatidylinositol is preferentially located in the cytosolic part of the membrane (inner leaflet). The negatively charged inositol head group can induce local reorganization through strong electrostatic interactions. Figure 6A shows the difference in the microdomain count between HB and ADB systems containing cholesterol, ceramide, sphingomyelin, and phosphatidylinositol of various sizes in the inner leaflet. In the GM region, the smaller microdomains (sizes 2 & 3) are abundant in healthy conditions, while higher-order microdomains are abundant in diseased conditions. In the WM region, the smaller microdomains (sizes 2, 3 & 4) are abundant in ADB systems, while higher-order domains are abundant in the HB system; however, the number of higher-order aggregates is much lower in WM than in the GM region of the brain. Figure 6B shows the schematic representation of POPI, PSM, CER, and CHL in the inner leaflet in different studied conditions, with all other membrane lipids shown as sticks. Clearly, the spatiotemporal organization of microdomains containing cholesterol, ceramide, sphingomyelin, and phosphatidylinositol is visibly different in healthy and diseased model membranes of GM regions. However, the distributions of microdomains are less apparent in healthy and diseased membrane models of the WM region.

**Figure 6:**
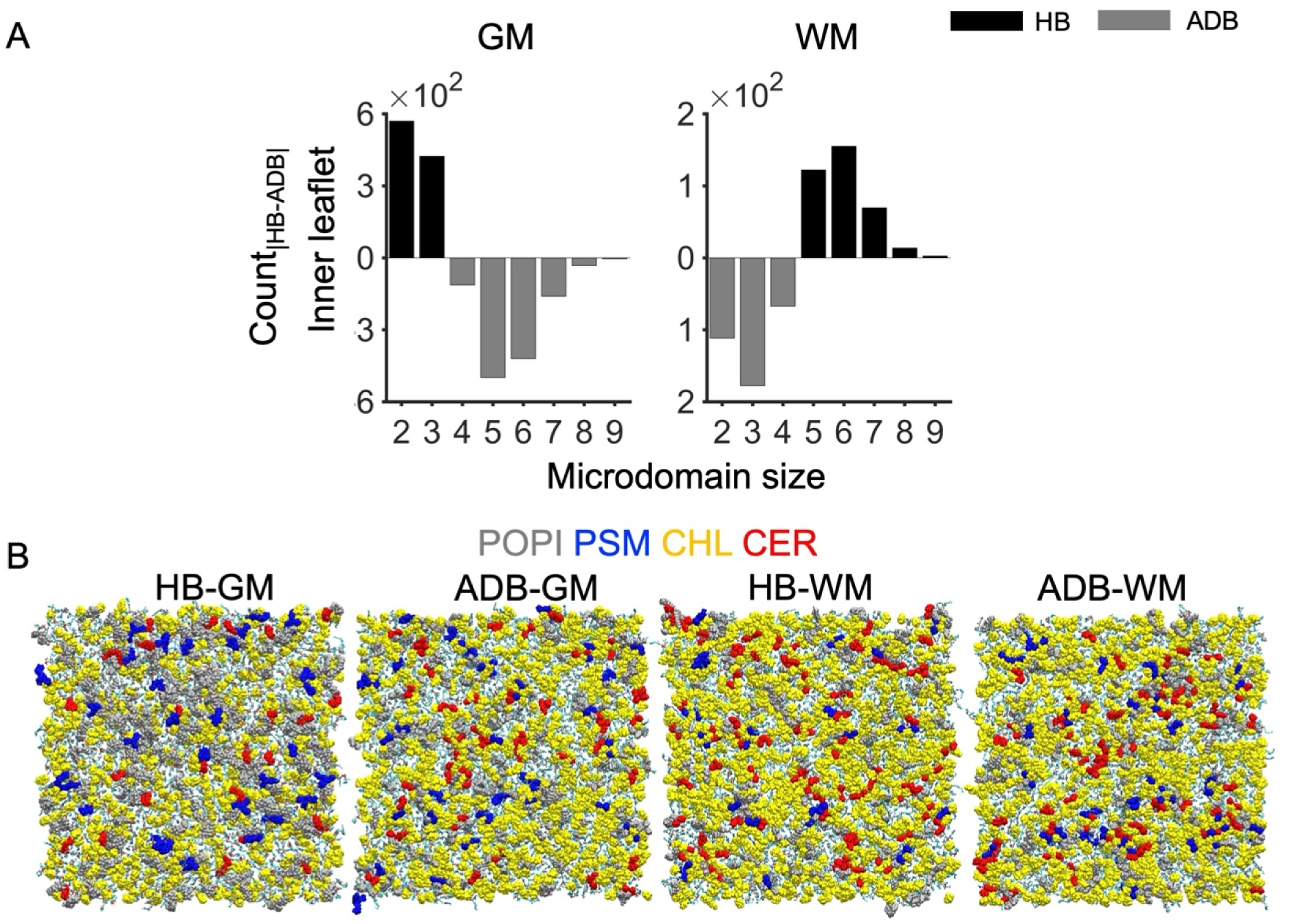
A) Difference in the number of aggregates containing CER, CHL, POPI, and PSM in the inner leaflets of healthy and diseased conditions of the GM and WM region, normalized with respect to the number of cholesterols. B) Schematic representation of a representative inner leaflet showing the spatial distribution of POPI, CER, PSM, and CHL among other components in the GM and WM region containing healthy and diseased membranes. (HB: healthy membrane, ADB: AD-mimicking diseased model membrane, GM: gray matter, WM: white matter).

## Conclusions

AD-associated changes in the lipid composition of GM and WM regions of the brain alter the structural and spatial organization of neuronal plasma membranes. Alterations, particularly in the structural features of the membrane, such as thickness and microdomain distribution, indicate the impact of lipid compositions on the neuronal membrane homeostasis and function in diseased conditions. Computational lipidomics of model membranes, developed from the postmortem of frozen brain tissues of subjects with AD pathology, namely Braak stage V-VI, and moderate-severe vascular pathology, and age-matched controls, reveals that the cholesterol-ceramide-sphingomyelin-enriched domains are abundant in the diseased membrane, particularly in the gray matter region. We have observed that, within this group of participants, membrane lipids of gray matter are more perturbed, both compositionally and structurally, than those of white matter. Detailed analysis of individual microdomains revealed that cholesterol-exclusive and cholesterol-enriched microdomains are abundant and more stable in the diseased membrane of the gray matter. We have also modelled bilayer asymmetry in our developed healthy and diseased model. Phosphatidylinositol is preferentially located in the cytosolic part of the membrane (inner leaflet). Our data show differences in the microdomains count between healthy and diseased membranes, with cholesterol, ceramide, sphingomyelin, and phosphatidylinositol, of various sizes in the inner leaflet. Particularly, the higher-order microdomains are abundant in the diseased membranes in the gray matter.

## Associated Content

## Supporting Information

Details of leaflet-wise composition, number density distribution of lipids across leaflets, membrane structural properties, radial distribution function profiles of CER (O3) and PSM (O3) relative to CHL (O3) in GM and WM of healthy and AD membrane models, lipid enrichment distribution profiles, residence time profile of microdomains sizes varying from 3 to 8.

## Abbreviations

CHL: Cholesterol
POPC: 1-palmitoyl-2-oleoyl-*sn*-glycero-3-phosphocholine
POPE: 1-palmitoyl-2-oleoyl-*sn*-glycero-3-phosphoethanolamine
POPS: 1-palmitoyl-2-oleoyl-*sn*-glycero-3-phosphoserine
POPI: 1-palmitoyl-2-oleoyl-*sn*-glycero-3-phosphoinositol
PSM: *N*-palmitoyl-d-erythro-sphingosine phosphorylcholine
CER: *N*-palmitoyl-d-erythro-sphingosine
POPA: 1-palmitoyl-2-oleoyl-sn-glycero-3-phosphatidic acid
POPG: 1-Palmitoyl-2-oleoylphosphatidylglycerol

## Author Information

### Notes

The authors declare no competing financial interest.

## Acknowledgments

SC gratefully acknowledges AMD for providing high-performance computing support to conduct research through the AMD AI & HPC Fund Research Cluster program. SP acknowledges the Department of Biotechnology for the Research Associate fellowship.

