## Supporting Information for "Modeling the organizational heterogeneity of lipid-enriched microdomains in the neuronal membranes of gray and white matter of Alzheimer’s brain: A computational lipidomics study"

**Table S1**: The composition of individual lipid components in outer and inner leaflets across the four systems studied. (OL: outer leaflet, IL: inner leaflet).

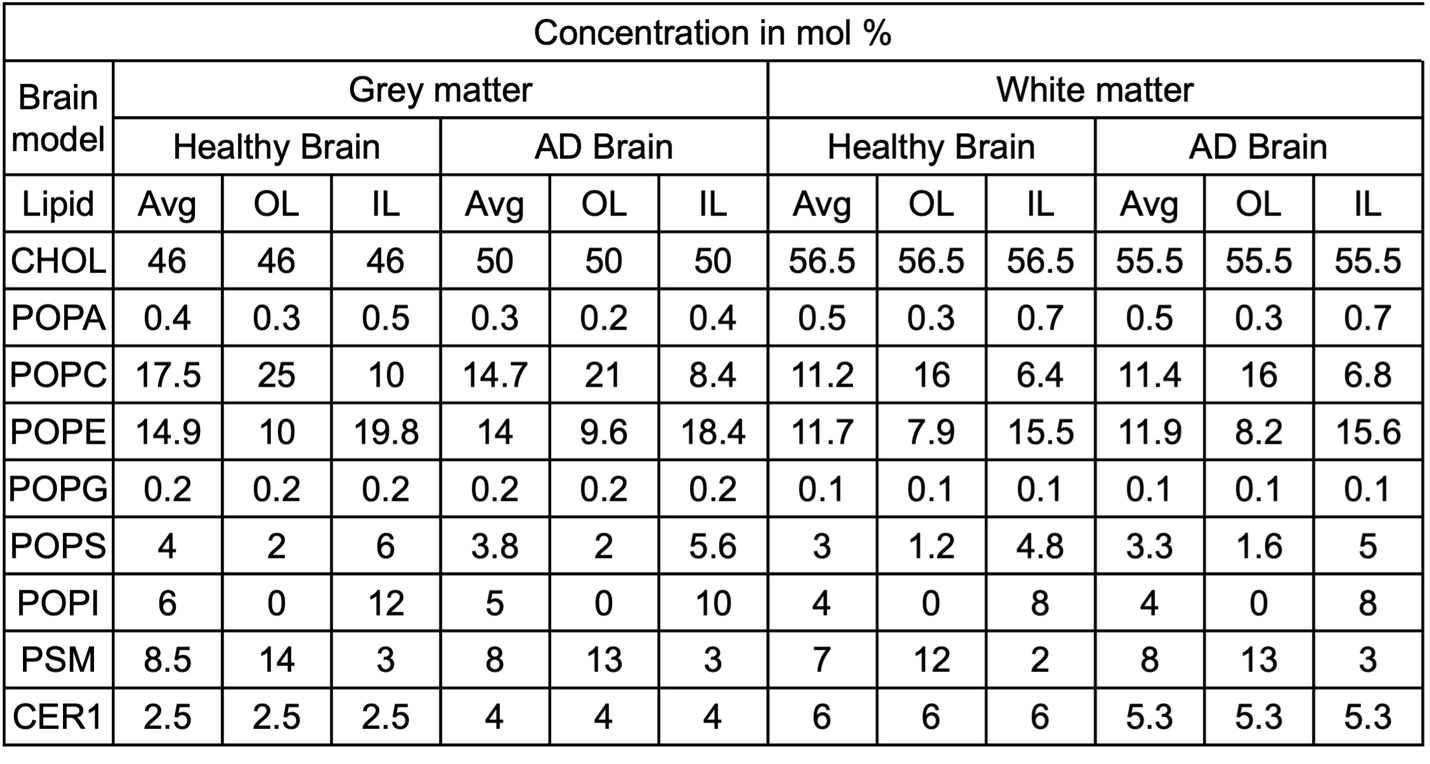

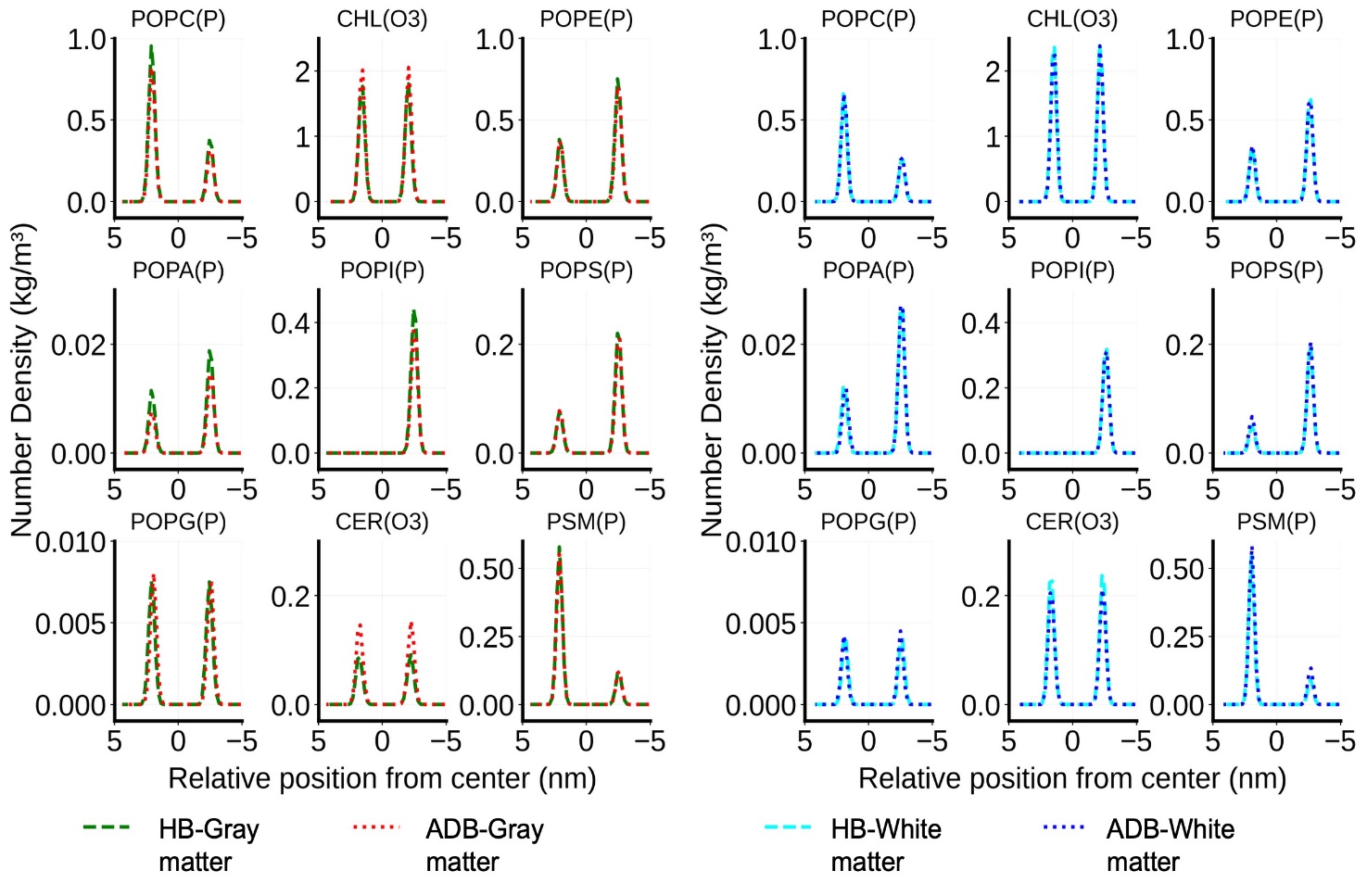

**Figure S1**: The number density distribution of all lipids across the four systems. HB: healthy membrane model, ADB: AD-mimicking diseased model membrane.

**Table S2**: Membrane bilayer structural properties of all the systems. The area per lipid as determined using the “Phosphorous” atom of POPC, POPE, POPS, POPI, POPA, POPG, PSM, and the O3 atom of CER, CHL as reference. The membrane thickness is determined using the “Phosphorous” atom of POPC, POPE, POPS, POPI, POPA, POPG, and PSM as the reference.

| **Property** | **Gray matter** | | **White matter** | |
| --- | --- | --- | --- | --- |
|  | Healthy Brain | AD Brain | Healthy Brain | AD Brain |
| Area per lipid  (Phospholipids- ‘P’, CER- ‘O3’, CHL- ‘O3’) (nm^2^) | 0.42 ± 0.0014 | 0.42 ± 0.0013 | 0.41 ± 0.0012 | 0.41 ± 0.0012 |
| Thickness (Phospholipids- ‘P’) (nm) | 4.59 ± 0.0128 | 4.57 ± 0.0125 | 4.53 ± 0.0136 | 4.54 ± 0.0130 |
| Interdigitation | 0.208 ± 0.021 | 0.205 ± 0.020 | 0.201 ± 0.187 | 0.201 ± 0.018 |
| Cholesterol Tilt angle  (‘C13-C10’) | 12.05 ± 0.326 | 11.75 ± 0.314 | 11.24 ± 0.289 | 11.27 ±0.285 |

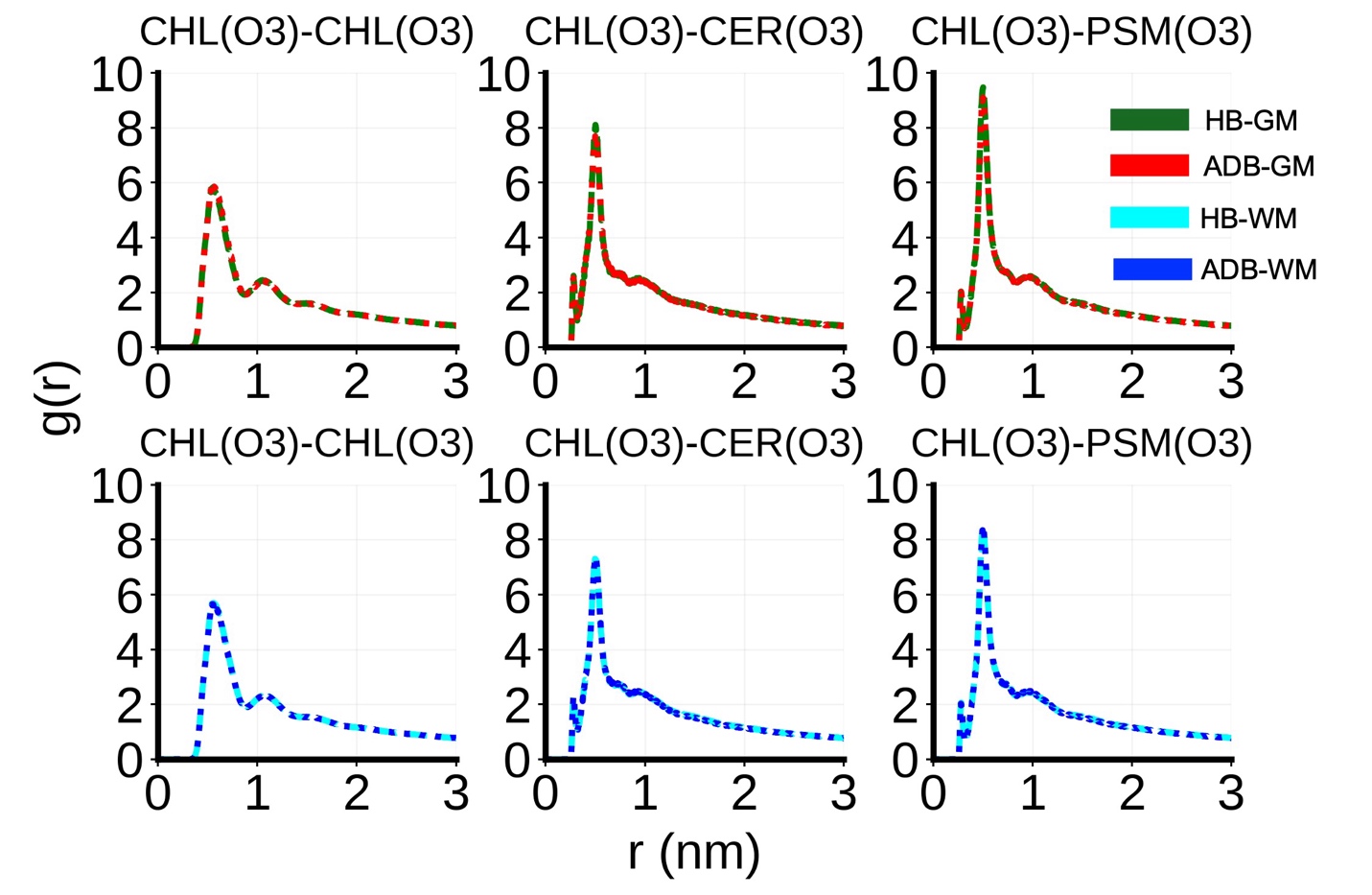

**Figure S2:** The radial density distribution profile of cholesterol, ceramide, and sphingomyelin around cholesterol. (HB: healthy membrane model, ADB: AD-mimicking diseased model membrane, GM: gray matter, WM: white matter).

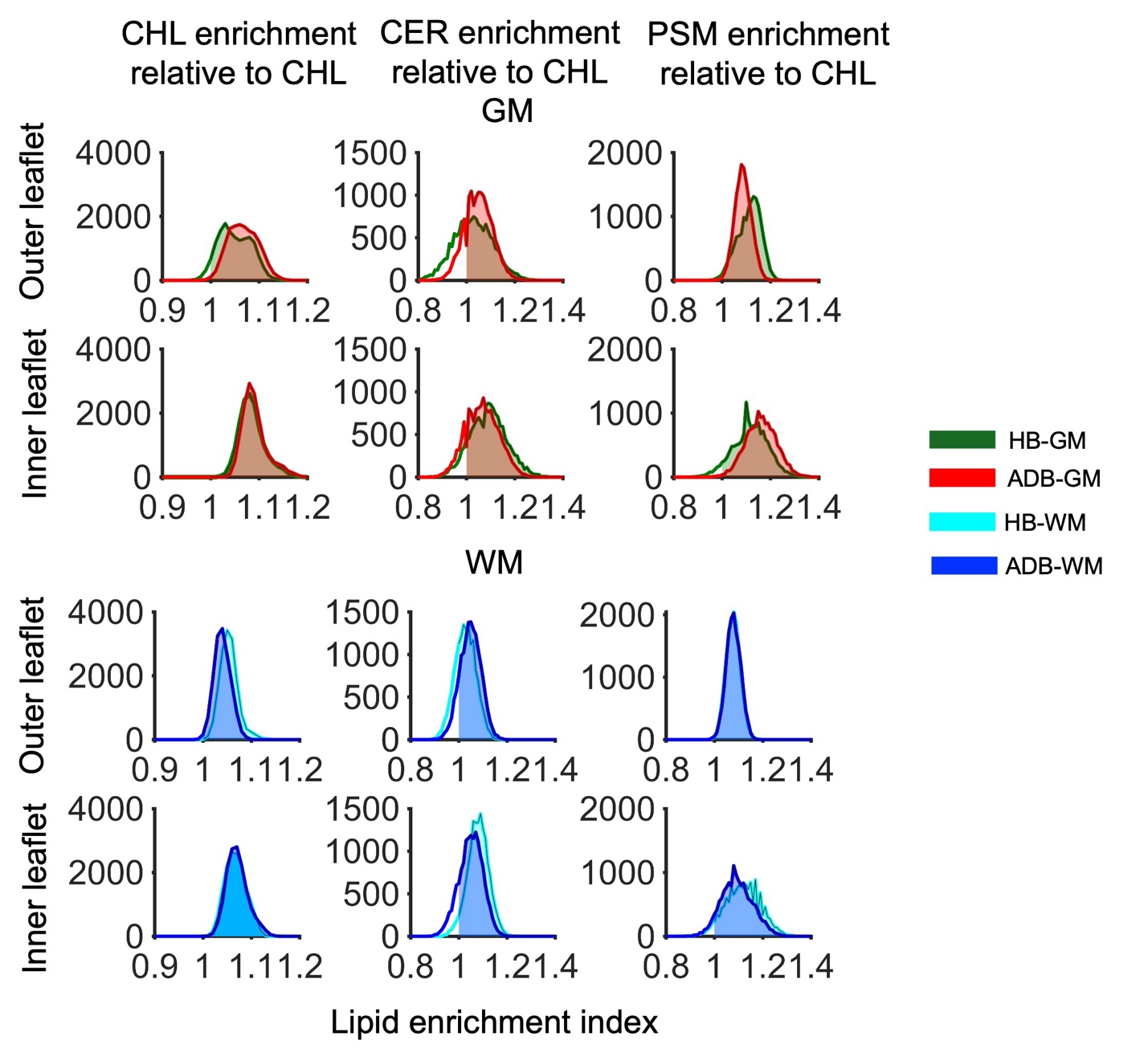

**Figure S3**: The distribution of enrichment index of cholesterol, ceramide, and sphingomyelin relative to cholesterol across the leaflets of healthy (HB) and diseased membranes (ADB) in the GM (gray matter) and WM (white matter) region. (HB: healthy membrane model, ADB: AD-mimicking diseased model membrane, GM: gray matter, WM: white matter).

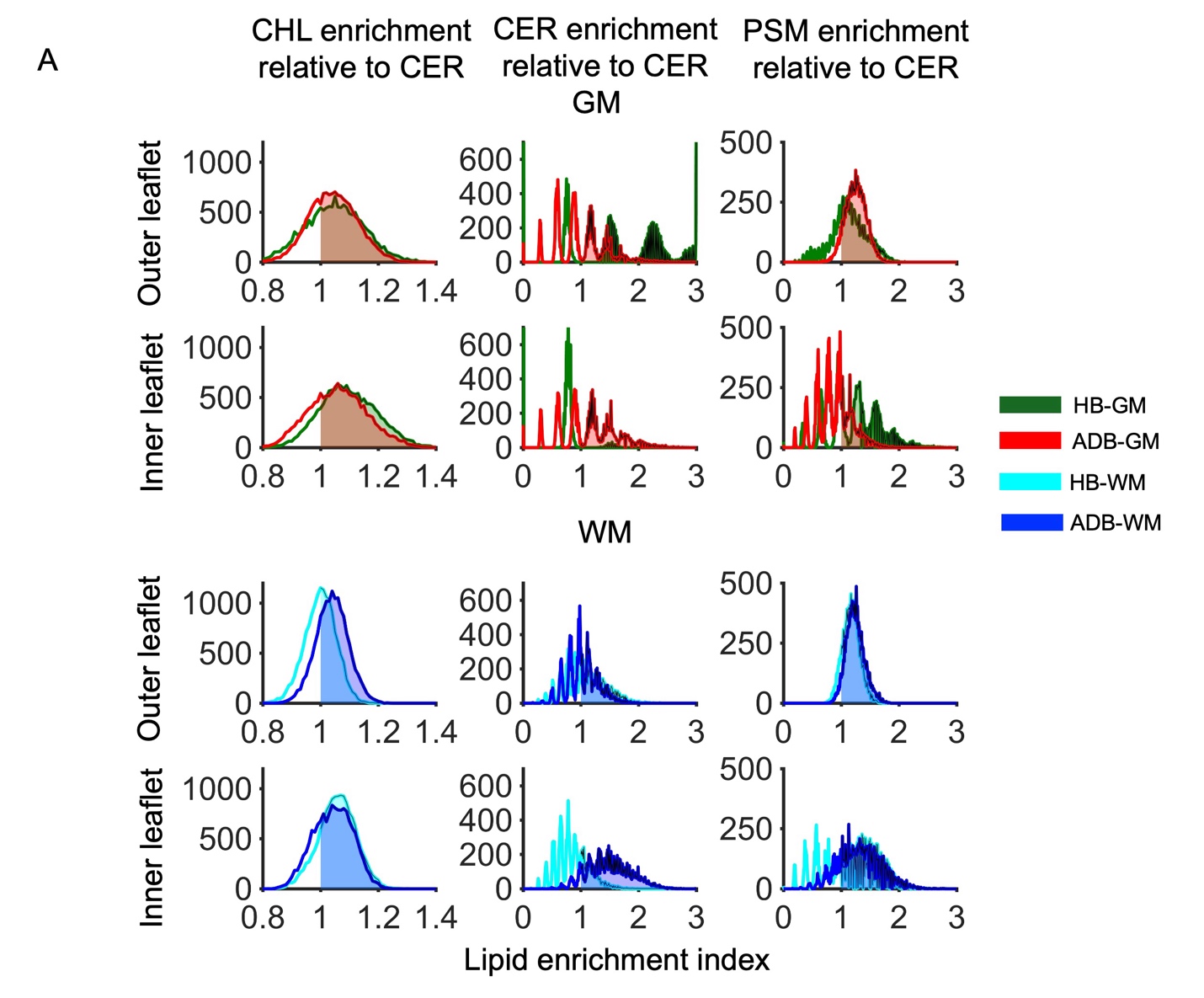

**Figure S4**: The enrichment index distribution of cholesterol, ceramide, and sphingomyelin relative to ceramide across the leaflets of healthy (HB) and diseased membranes (ADB) in the GM (gray matter) and WM (white matter) region. (HB: healthy membrane model, ADB: AD-mimicking diseased model membrane, GM: gray matter, WM: white matter).

**Table S3**: Area under the curve values, obtained from the enrichment index value distribution curve (Figure S3), for regions with enrichment values greater than or equal to 1. (HB: healthy membrane model, ADB: AD-mimicking diseased model membrane, OL: outer leaflet, IL: inner leaflet, GM: gray matter, WM: white matter, CHL-Cholesterol, CER- *N*-palmitoyl-d-erythro-sphingosine, PSM- *N*-palmitoyl-d-erythro-sphingosine phosphorylcholine).

| **Membrane type** | **CHL-CHL** | | **CER-CHL** | | **PSM-CHL** | |
| --- | --- | --- | --- | --- | --- | --- |
|  | OL | IL | OL | IL | OL | IL |
| **HB-GM** | 141.85 | 150.03 | 94.29 | 132.43 | 148.27 | 141.21 |
| **ADB-GM** | 149.25 | 150.03 | 120.71 | 123.96 | 149.48 | 149.18 |
| **HB-WM** | 149.98 | 150.02 | 109.19 | 143.88 | 149.5 | 142.4 |
| **ADB-WM** | 149.07 | 150.02 | 129.07 | 125.61 | 149.5 | 139.7 |

**Table S4**: Area under the curve values, obtained from the enrichment index value distribution curve (Figure S4), for regions with enrichment values greater than or equal to 1. (HB: healthy membrane model, ADB: AD-mimicking diseased neuronal model membrane, OL: outer leaflet, IL: inner leaflet, GM: gray matter, WM: white matter, CHL-Cholesterol, CER- *N*-palmitoyl-d-erythro-sphingosine, PSM- *N*-palmitoyl-d-erythro-sphingosine phosphorylcholine).

| **Membrane type** | **CHL-CER** | | **CER-CER** | | **PSM-CER** | |
| --- | --- | --- | --- | --- | --- | --- |
|  | OL | IL | OL | IL | OL | IL |
| **HB-GM** | 101.37 | 125.61 | 86.5 | 2.9 | 107.91 | 100.8 |
| **ADB-GM** | 103.36 | 107.54 | 64.7 | 85.5 | 138.12 | 50.8 |
| **HB-WM** | 76.51 | 122.76 | 87.1 | 45.2 | 130.98 | 92.53 |
| **ADB-WM** | 114.53 | 108.13 | 77.2 | 136.3 | 142.34 | 121.62 |

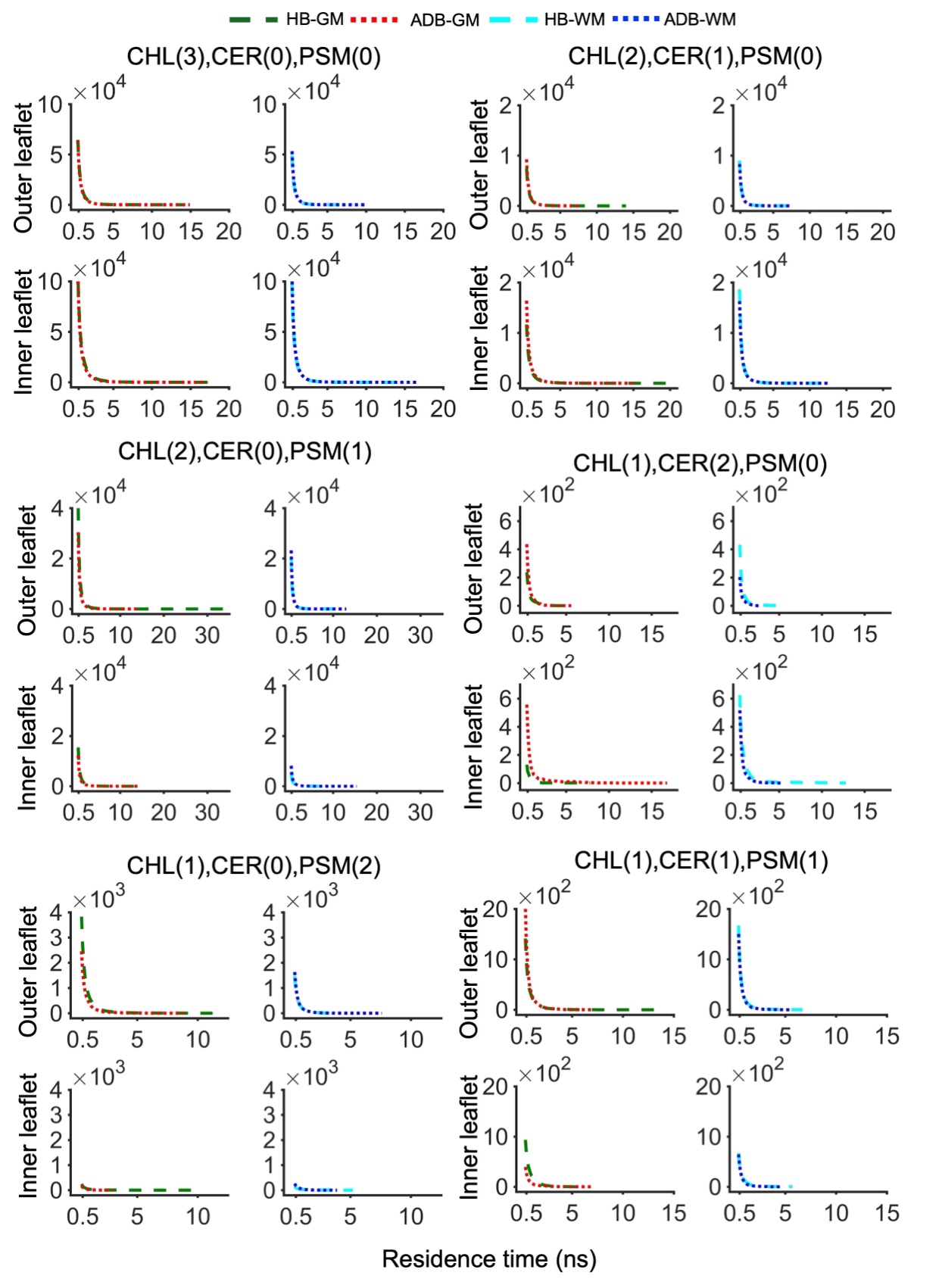

**Figure S5**: Residence time distributions of different compositions of microdomains of size 3 calculated from the 1.5 µs concatenated trajectory. (HB: healthy membrane model, ADB: AD-mimicking diseased model membrane, GM: gray matter, WM: white matter).

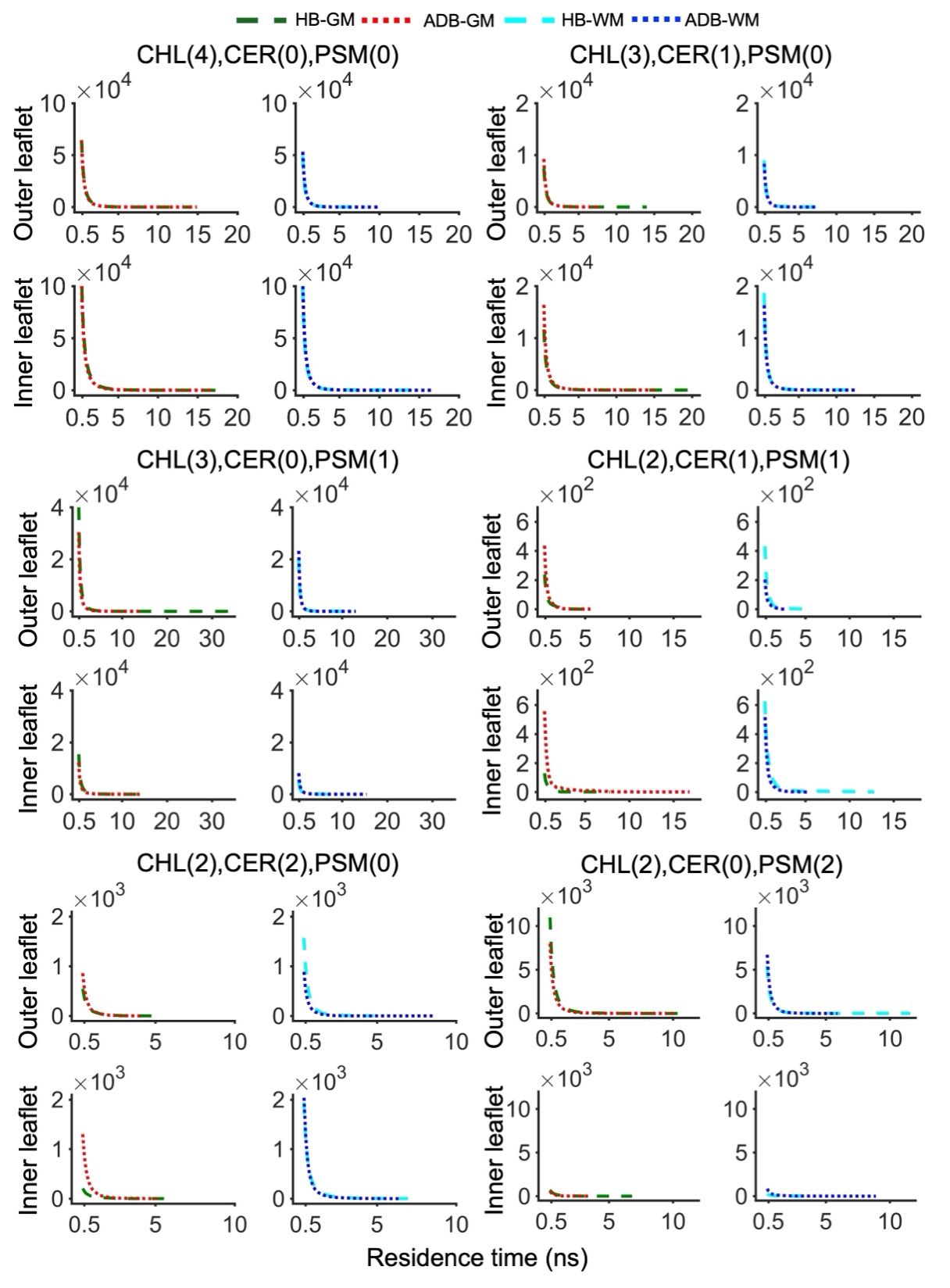

**Figure S6**: Residence time distributions of different compositions of microdomains of size 4 calculated from the 1.5 µs concatenated trajectory. (HB: healthy membrane model, ADB: AD-mimicking diseased model membrane, GM: gray matter, WM: white matter).

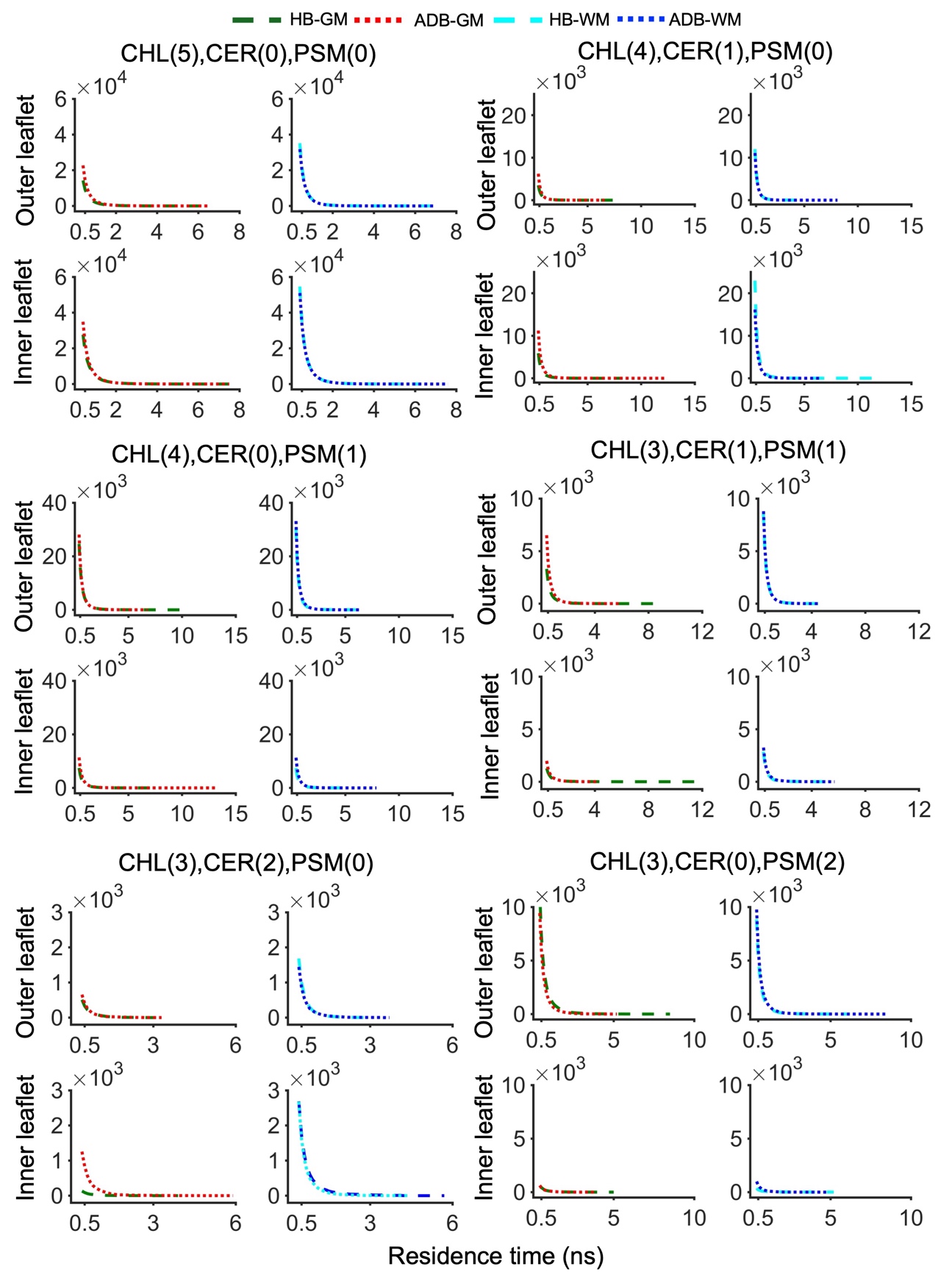

**Figure S7:** Residence time distributions of different compositions of microdomains of size 5 calculated from the 1.5 µs concatenated trajectory. (HB: healthy membrane model, ADB: AD-mimicking diseased model membrane, GM: gray matter, WM: white matter).

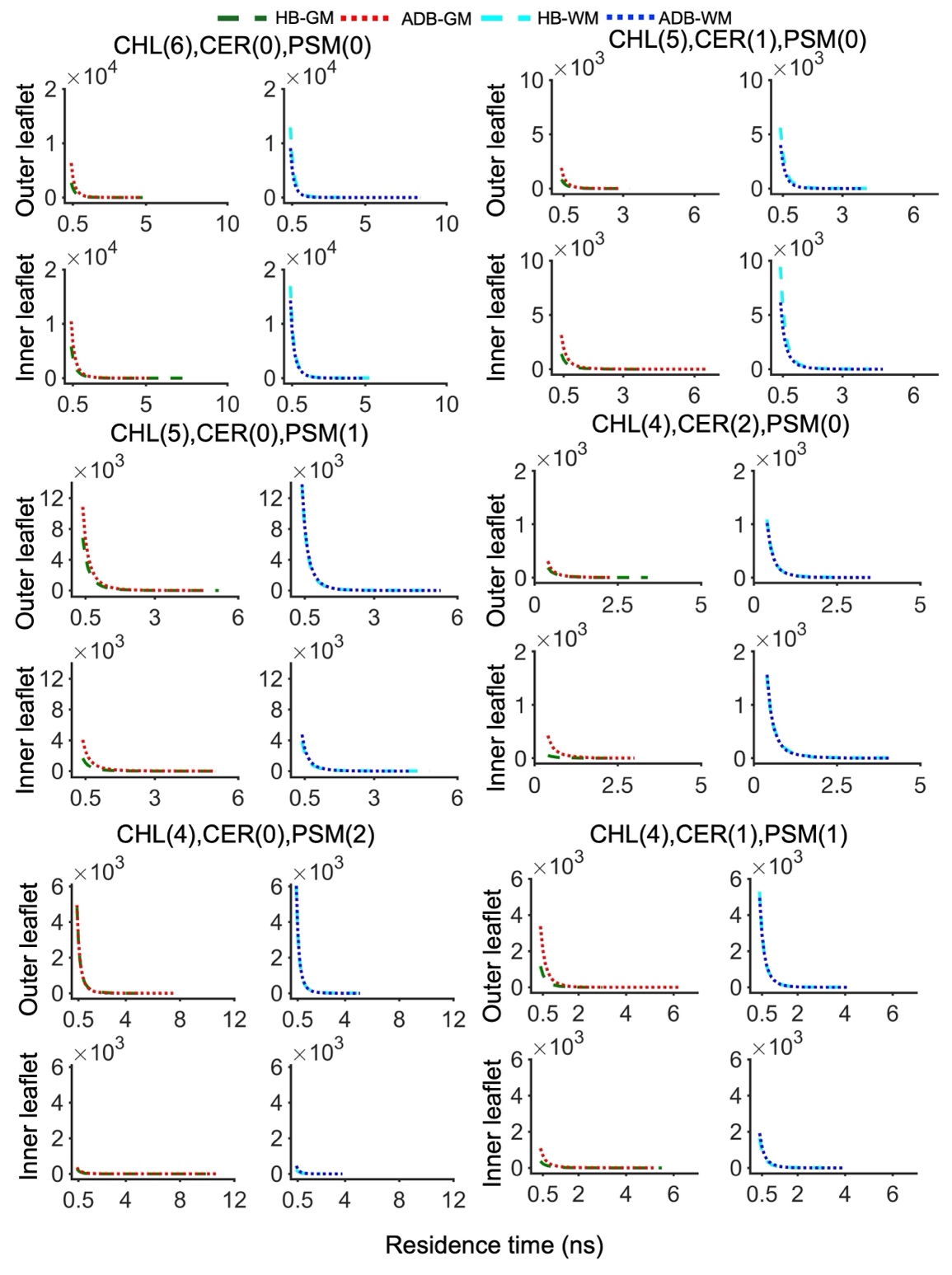

**Figure S8:** Residence time distributions of different compositions of microdomains of size 6 calculated from the 1.5 µs concatenated trajectory. (HB: healthy membrane model, ADB: AD-mimicking diseased model membrane, GM: gray matter, WM: white matter).

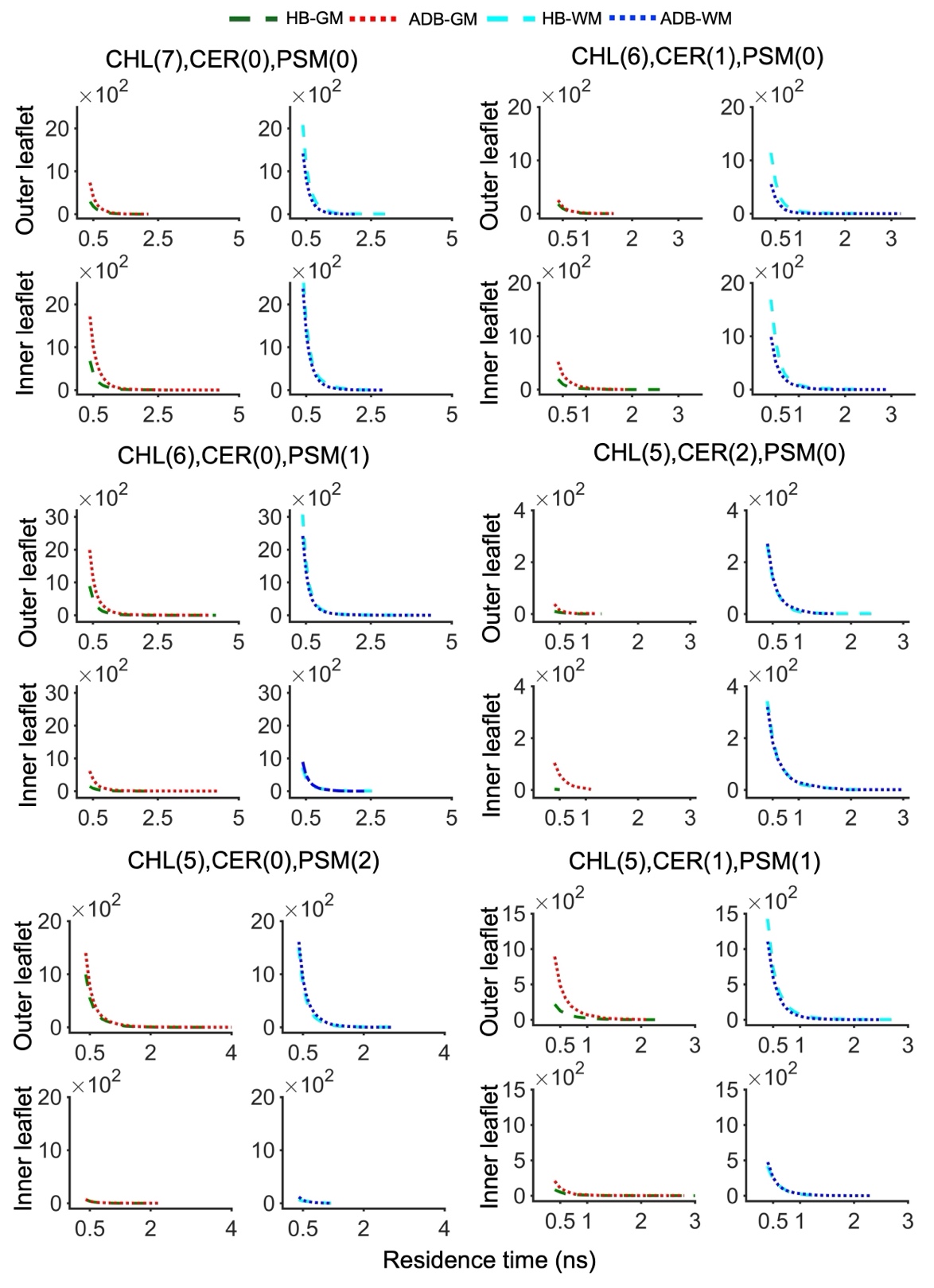

**Figure S9**: Residence time distributions of different compositions of microdomains of size 7 calculated from the 1.5 µs concatenated trajectory. (HB: healthy membrane model, ADB: AD-mimicking diseased model membrane, GM: gray matter, WM: white matter).

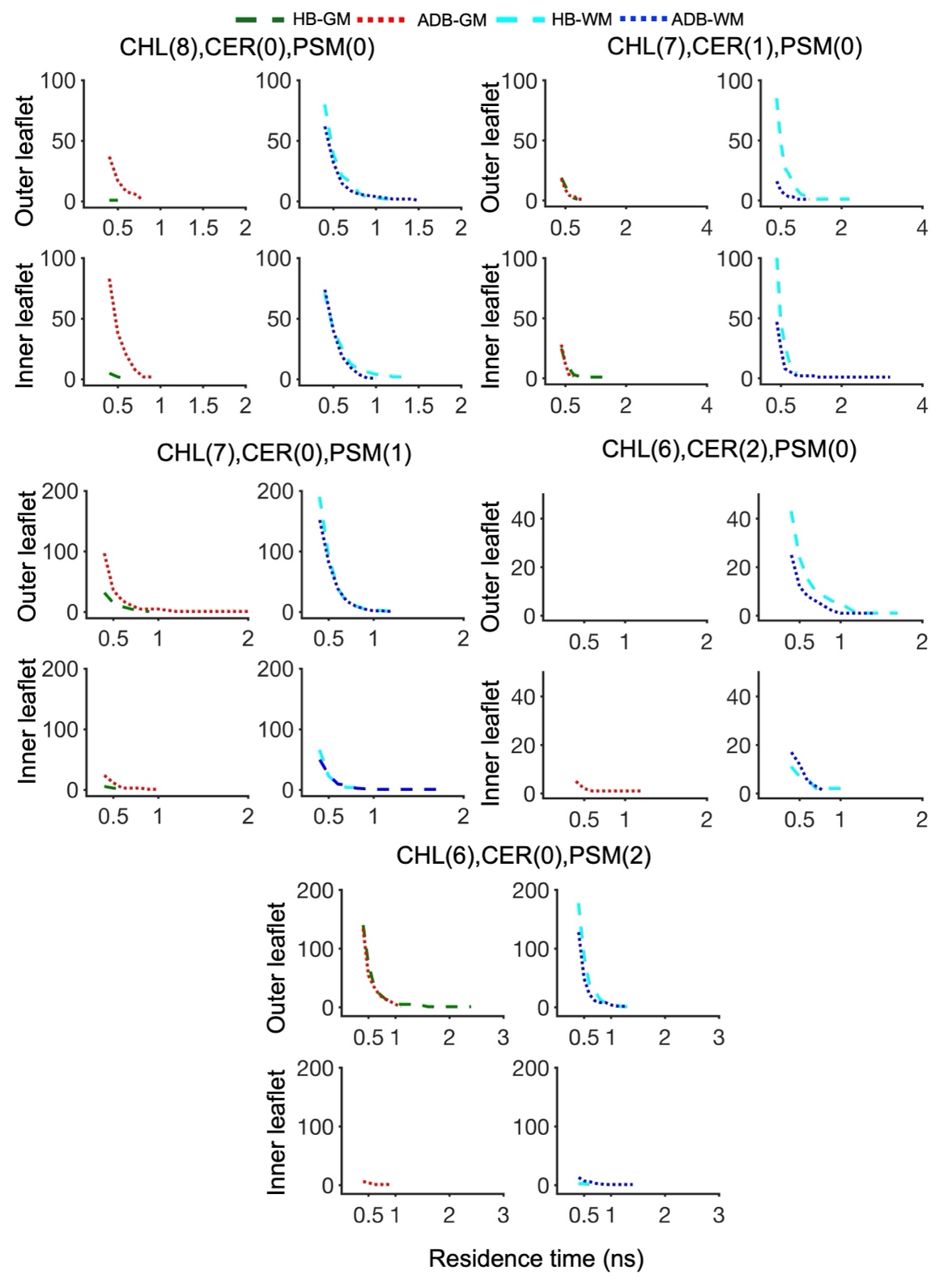

**Figure S10:** Residence time distributions of different compositions of microdomains of size 8 calculated from the 1.5 µs concatenated trajectory. (HB: healthy membrane model, ADB: AD-mimicking diseased model membrane, GM: gray matter, WM: white matter).
